# Selective Attention Modulates Neural Envelope Tracking of Informationally Masked Speech in Healthy Older Adults

**DOI:** 10.1101/2020.10.13.337378

**Authors:** Ira Kurthen, Jolanda Galbier, Laura Jagoda, Pia Neuschwander, Nathalie Giroud, Martin Meyer

## Abstract

Speech understanding in noisy situations is compromised in old age. This study investigated the energetic and informational masking components of multi-talker babble noise and their influence on neural tracking of the speech envelope in a sample of healthy older adults. Twenty-three older adults (age range 65 - 80 years) listened to an audiobook embedded in noise while their electroencephalogram (EEG) was recorded. Energetic masking was manipulated by varying the signal-to-noise ratio (SNR) between target speech and background talkers and informational masking was manipulated by varying the number of background talkers. Neural envelope tracking was measured by calculating temporal response functions (TRFs) between speech envelope and EEG. Number of background talkers, but not SNR modulated the amplitude of an earlier (around 50 ms time lag) and a later (around 300 ms time lag) peak in the TRFs. Selective attention, but not working memory or peripheral hearing additionally modulated the amplitude of the later TRF peak. Finally, amplitude of the later TRF peak was positively related to accuracy in the comprehension task. The results suggest that stronger envelope tracking is beneficial for speech-in-noise understanding and that selective attention is an important ability supporting speech-in-noise understanding in multi-talker scenes.

## Introduction

Older adults often report difficulties in understanding speech in background noise (CHABA, 1988; Pichora-Fuller, 1995; Humes et al., 2012), even when they are considered normal hearing based on pure-tone thresholds (Füllgrabe et al., 2015). Speech understanding difficulties are augmented in the presence of age-related hearing loss (ARHL) (Kortlang et al., 2016). ARHL, or presbycusis, is the most common form of sensorineural hearing loss, and it is one of the most prevalent age-related conditions, estimated at approximately 20% at age 60, 50% at age 70 and 70% to 80% at age 80 and older (Goman and Lin, 2016; Bisgaard and Ruf, 2017). It can result in multiple unwanted outcomes like social isolation (Weinstein and Ventry, 1982; Mick et al., 2014; Ciorba et al., 2012) and potentially cognitive decline (Maharani et al., 2019) and eventually dementia (Lin et al., 2011). It is assumed that the higher risk for social isolation stems at least partly from the background noise typically associated with social situations. Currently, hearing aids constitute the only evidence-based treatment for age-related hearing loss. While they can improve the quality of life for affected individuals (Stark and Hickson, 2004; Johnson et al., 2018), the disturbance of noisy situations is still the most common complaint after the fitting of a hearing aid (Bertoli et al., 2009). Therefore, understanding the processes and abilities that lead to successful speech-in-noise understanding in older adults is key to helping develop additional treatments for ARHL which let older adults maintain their social relationships.

Indeed, one of the most difficult communication situations is listening to a single speaker in the presence of other talkers, which is known as the “cocktail party problem” (Cherry, 1953). Both the target signal (the speech signal the listener aims to attend to) and the noise (the speech signals of other talkers which the listener is trying to ignore) are speech signals. Hence, the frequency bands in which these signals contain energy will tend to overlap, a phenomenon which is commonly referred to as “energetic masking” (EM, Brungart, 2001). However, speech-on-speech masking presents an additional challenge that cannot be explained only by an overlap in energy frequency bands. This additional type of masking has been labelled “informational masking” (IM, Brungart, 2001), and it is notoriously difficult to define, the common ground of all definitions being that its masking properties are nonenergetic, i.e. not explained by overlap in energy frequency bands (Durlach et al., 2003; Shinn-Cunningham, 2008; Rosen et al., 2013).

The amount of IM depends on the similarity of target and distractor, and, consequently, it can be reduced by increasing the dissimilarity between target and background talkers, for example by moving the distractor talker to a different location (Kidd et al., 1998) or by introducing sex differences between target and distracting talkers (Brungart, 2001). IM has also been shown to take more effect when the background talkers speak the same language as the target speaker than when they speak a foreign language (Rhebergen et al., 2005; Van Engen and Bradlow, 2007; Garcia Lecumberri and Cooke, 2006; Brouwer et al., 2012). Additionally, IM increases when the distractor language is known to the listeners, compared to an unknown language (Garcia Lecumberri and Cooke, 2006). These results have been explained on the basis of an increased cognitive load because of language-decoding mechanisms that take place when a known language is presented as distractor (Cooke et al., 2008). This conclusion has been strengthened by the finding that IM is stronger when distractor speech consists of meaningful as opposed to semantically anomalous sentences (Brouwer et al., 2012).

According to Shinn-Cunningham (2008), IM may be related to failures of object-based attention, either because of failures in object formation, which occur when separate sources in a scene cannot be separated from one another, or because of failures in object selection, which occur when top-down attention is directed to the distractor rather than the target. Object formation may be more difficult for hearing-impaired individuals, because of a spectrotemporally degraded representation of the speech input (Shinn-Cunningham and Best, 2008). This degraded representation results in perceptually more similar target and distractor objects. This similarity, in turn, leads to more difficulties in object selection (Shinn-Cunningham and Best, 2008), which can explain speech understanding difficulties of hearing-impaired individuals in multi-talker scenes. The Framework for Understanding Effortful Listening (Pichora-Fuller et al., 2016) comprehensively describes how attention governs the allocation of cognitive capacity to cope with difficult listening situations.

Another cognitive ability that is relevant for speech-in-noise understanding is working memory. Because masked speech as well as peripheral hearing loss can result in a degraded representation of the input signal, the role of working memory in phonological and lexical retrieval and in pattern matching as posited by the Ease of Language Understanding model (Rönnberg et al., 2013) is significant in a multi-talker scene. Indeed, especially for older individuals, working memory capacity is a very reliable predictor for speech in noise understanding (Zekveld et al., 2013; Moore et al., 2014; Besser et al., 2013), although see some counterevidence (Schoof and Rosen, 2014).

The “resource allocation hypothesis “ or “information degradation hypothesis “ (Wayne and Johnsrude, 2015; Nixon et al., 2019) underscores the importance of working memory capacity as a pool of resources that can be spent either on processing the sensory input or performing higher-level computations on that input. The related “effortfulness hypothesis” (e.g. McCoy et al., 2005) poses that under adverse listening conditions, resources are spent on processing of the challenging stimuli, which may later not be available for performing mental computations on the input, like encoding semantic content into memory. To measure performance in both of these domains (perceptual processing and semantic encoding) in our study, our participants had to complete two tasks: an *intelligibility task*, which simply tested how well participants could follow the target speaker, and a *comprehension task*, which tested participants’ memory of the lexical content of the target speech signal. If there are no differences in intelligibility, there might be differences in comprehension, depending on how many resources were spent during the earlier task.

### Age-related Changes in Acoustic Cue Processing and Neural Envelope Tracking

With speech being an acoustic signal, the way the processing of acoustic cues changes with age is also of considerable importance. In general, aging is accompanied by a slowing of many processes (Salthouse, 1996, 2000), and this does not seem to be different in the acoustic domain. The acoustic signature of speech can generally be divided into two parts: rapidly changing acoustic cues, the temporal fine structure, and slowly changing acoustic cues, the temporal envelope (e.g. Drullman, 1995; Shannon et al., 1995; Smith et al., 2002). Especially slowly changing envelope cues are crucial for successful speech understanding (Shannon et al., 1995; Liem et al., 2014). A number of studies have shown that while aging is accompanied by decrease in the ability to process temporal fine structure, the ability to process slowly changing cues, like the temporal envelope, is preserved (Gordon-Salant and Fitzgibbons, 1993; Schneider and Pichora-Fuller, 2001; Wingfield et al., 2000, 1992; Lorenzi et al., 2006; Schneider et al., 2010; Meyer et al., 2018; Giroud et al., 2019). In conclusion, it appears that slow acoustic features of speech are an important resource for older adults to draw upon when understanding speech, especially in challenging listening situations.

On a neural level, it is presumed that initial encoding of speech occurs by means of entrainment of ongoing cortical oscillations to the speech envelope (e.g. Giraud and Poeppel, 2012; Gross et al., 2013). This entrainment to the envelope is also called “envelope tracking”, and studies with transcranial alternating current stimulation have provided evidence that it serves a causal role for (i.e., functionally contributes to) speech intelligibility (Riecke et al., 2018; Wilsch et al., 2018; Zoefel et al., 2018). Envelope tracking is a robust phenomenon, demonstrated by two studies that demonstrated envelope tracking even in severe acoustic interference from a competing talker (SNR between attended and ignored talker up to −8 dB, Ding and Simon, 2012, 2013). Nevertheless, it exhibits considerable inter-individual variability (Lam et al., 2018). When comparing envelope tracking younger and older adults, older adults on average typically show a stronger cortical response than younger adults Presacco et al. (2016b); Decruy et al. (2019). Similarly, a neural over-representation of the envelope compared with the temporal fine structure has been demonstrated in individuals with ARHL (Anderson et al., 2013b), probably because ARHL mainly occurs in higher frequencies, which leaves the envelope relatively intact. It is currently unclear whether higher envelope tracking is adaptive and reflects compensatory mechanisms or whether it constitutes a true “over”-representation, which can hinder processing of the temporal fine structure (Decruy et al., 2019). At least in a study with young adults, stronger envelope tracking was highly positively correlated with subjectively rated speech-in-noise intelligibility (Ding and Simon, 2013).

Envelope tracking is especially relevant in a cocktail-party environment. The study by O’Sullivan et al. (2015) showed that envelope tracking predicted target speech intelligibility in a cocktail-party situation. A study by Vander Ghinst et al. (2016) investigated how envelope tracking and noise level of multi-talker babble noise level was related. They found that as the noise level increased, envelope tracking decreased. Selective attention, which is an important ability for auditory object formation and selection (Shinn-Cunningham, 2008), is also important for envelope tracking. In a study by Kerlin et al. (2010), selective attention was shown to increase the gain of ongoing speech representations in a cocktail party scenario. Zion Golumbic et al. (2013) showed that speech representations in cocktail party scenarios become more and more sharpened to the target speech while a sentence unfolds and while the signal progresses through the processing hierarchy. Entrainment of cortical oscillations to the speech envelope may even serve to suppress the competing speech signal (Horton et al., 2013), which is an ability that seems to be impeded with higher levels of peripheral hearing loss (Petersen et al., 2017). Taken together, these results suggest that envelope tracking is a necessary step in speech processing, that envelope tracking can be increased by exerting selective attention, and that stronger envelope tracking is positively related with speech intelligibility in multi-talker situations.

Although working memory is a reliable predictor of speech-in-noise understanding in behavioral studies, a recent study found only weak evidence for an involvement of working memory in speech envelope tracking (Decruy et al., 2019). However, that particular study and many others have measured envelope tracking by performing envelope reconstruction from observed brain activity using backwards modeling. This method provides a quantification of envelope reconstruction fidelity, but it does not take into account temporal aspects of envelope tracking. In our study, we measured envelope tracking by fitting TRFs (also called auditory-evoked spread spectrum analyses (AESPAs), (Lalor et al., 2009), which constitute a form of forward modeling that allows to observe how the envelope tracking unfolds across time. One can therefore specifically analyze envelope tracking in time windows during which cognition is known to influence brain activity (O’Sullivan et al., 2015). Power et al. (2012) have used this method to detect a attention-modulated peak in the AESPA at a lag of around 200 ms.

### Study Design and Hypotheses

Reconciling the two main findings that cognitive ability and hearing loss are related to speech in noise understanding and that aging and selective attention modulate envelope tracking, it is useful to ask whether envelope tracking is a mechanism by which cognition exerts its positive influence on speech-in-noise understanding. Specifically, we aimed to ascertain whether the EM and IM components of multi-talker babble noise would affect envelope tracking, and if yes, whether they affect it independently or with additive effects.

We therefore designed this study to investigate envelope tracking in the context of EM and IM and to study the relationship between cognition and envelope tracking in multi-talker babble noise. To this end, while their electroencephalogram (EEG) was recorded, our participants listened to speech in multi-talker babble noise that was designed within a 2×2 design: we manipulated the SNR between target speaker and background noise for EM and the number of background talkers (nBT) for IM, as it has been shown that IM arising from lexical interference decreases when nBT increases (Rosen et al., 2013).

Because the addition of other talkers can also influence EM (Rosen et al., 2013), and because the effectiveness of IM strongly depends on similarity between talker and distractor (e.g. sex, semantic content, spatial orientation), we decided to model the background noise from the same speaker who uttered the target signal. Additionally, EM in multi-talker babble noise strongly depends on whether the background talkers are currently speaking or pausing. To reduce the possibility for glimpsing, silent periods in the noise stimuli were trimmed. This way, we ensured that EM was always present and we can assume a monotonic increase in EM by lowering the SNR and a monotonic decrease in IM by adding more BT.

For the nBT, we decided to present two conditions with two background talkers (2 BT), because this is the most difficult multi-talker babble condition (Rosen et al., 2013; Freyman et al., 2004) and two conditions with eight background talkers (8 BT), because then, the background talkers mask one another and lexical interference is reduced. Different SNRs were tested in a pilot study, from which the SNRs 0 and 2 resulted. An overview of the experimental conditions can be found in Table 1.

**Table 1.** Experimental Conditions

| SNR (EM) | Number of Background Talkers (IM) |  |
| --- | --- | --- |
|  | 2 | 8 |
| 0 | high EM, high IM | high EM, low IM |
| 2 | low EM, high IM | low EM, low IM |

*Note.* This table shows the four experimental conditions. Each condition is defined by high or low EM and IM.

To account for speech processing difficulties that would propagate beyond mere speaker tracking, (effortfulness hypothesis McCoy et al., 2005), participants had to complete two tasks: an intelligibility task and a comprehension task. If there are no differences in performance in the intelligibility task between conditions, there might be differences in performance in the comprehension task that show up because of different amounts of resources remaining, after more or less of them were spent on processing the input, but not working on it or storing it.

To extract neural envelope tracking from the EEG recordings, we fitted Temporal Response Functions (TRFs) to the envelope of the target signal with functions provided by the mTRF toolbox for MATLAB Crosse et al. (2016). TRFs are forward models of the “time course of the neural response evoked by a unit power increase of the stimulus” (Ding and Simon, 2012, p.11856), and they contain timing and spatial information of the neural encoding process (Ding and Simon, 2012). In fact, they are (and look) similar to auditory event-related potentials (ERPs, Lalor et al., 2009), although they only represent neural activity in reaction to a specific feature, in our case the envelope, and not the net sum of all activity that is time-locked to stimulus onset. While cross-correlating envelope and EEG (Zoefel and VanRullen, 2016; Petersen et al., 2017) is a similar approach to estimating the neural response to the envelope, TRFs are better suited because the speech envelope exhibits significant autocorrelation, which causes temporal smearing in the cross-correlation approach (Crosse et al., 2016). In the mTRF toolbox, fitting of TRFs is achieved via ridge regression, which is among the best regularization methods for TRFs (Wong et al., 2018).

In a magnetencephalography study featuring an attended-speech paradigm, Ding and Simon (2013) found that amplitude of an early neuromagnetic component “M50” of the TRF linearly decreased with increasing noise level, but the amplitude of a later component “M100” of the TRF was not affected by noise until it drastically decreased between −6 and −9 dB. Ding and Simon (2012) found that the M100 peak of the TRF was modulated by attention, whereas the M50 was not. Also, up to −8 dB SNR, there was no effect of SNR on peak amplitude. However, Petersen et al. (2017) found that differences in SNR resulted in differences in envelope tracking, with tracking of attended speech being stronger in lower noise levels than in higher noise levels. Petersen et al. (2017) presented speech in subject-specific SNRs, while Ding and Simon (2012) and Ding and Simon (2013) used absolute SNRs. We also used absolute SNRs because we aimed to exclude any possible interference effects of EM and IM (Rosen et al., 2013) that could emerge differentially with subject-specific SNRs.

While earlier ERPs are associated with perceptual processing of exogenous stimuli, later peaking ERPs indicate cognitive and endogenous processing, like for example the P3b (van Dinteren et al., 2014; Giroud et al., 2016). Because EM is mainly a perceptual interference, we hypothesized that differences in SNR would manifest at early time points. Because IM due to lexical interference should tap higher-order cognitive resources, we hypothesized differences between TRFs in response to two vs. eight background talkers during a later time window typically associated with cognitive processing.

Because selective attention has been shown to increase the gain of ongoing speech representations during multi-talker babble noise (Kerlin et al., 2010), we hypothesized that a measurement of participants’ ability to exert selective attention would predict envelope tracking. Exploratively, because working memory is the most commonly found predictor for speech-in-noise understanding, we also tested whether working memory would predict envelope tracking. Finally, because envelope tracking has been shown to be related to hearing thresholds (Petersen et al., 2017), we also tested whether hearing thresholds would predict envelope tracking.

## Materials and Methods

### Participants

The sample consisted of 23 older adults (mean age = 70.96 yrs, sd = 3.72 yrs, 14 females). One more participant was tested but excluded due to technical issues during EEG recording. All participants were right-handed as assessed by the Annett Hand Preference Questionnaire (Annett, 1970) and reported no psychiatric or neurological disorders. Their native language was Swiss German and they had not learned another language before their seventh year of age. They did not play music for more than six hours per week and they did not wear a hearing aid. Their hearing loss (pure-tone average; PTA) did not exceed 60 dB in the frequencies 500, 1000, 2000, and 4000 Hz, and the mean difference in hearing thresholds between the two ears did not exceed 20 dB. They passed a screening procedure, in which the exclusion criteria were tested via questionnaires and their hearing thresholds were measured with a MAICO ST-20. Additionally, they were administered the Montreal Cognitive Assessment (Nasreddine et al., 2005) and were invited to further participate in the study when they scored 26 points or more.

The ethics committee of the Canton of Zurich approved the study (application no. 2017-00284). Written informed consent was obtained from all participants. Participants were compensated for their participation.

### Cognitive Tests

Selective attention was measured with the Eriksen-Flanker Task (Eriksen and Eriksen, 1974), administered in a computer version modeled after the version reported in Stins et al. (2007) and available in the PEBL software, Version 0.14 Mueller and Piper (2014). Working memory was assessed with the Sentence Span task from the working memory capacity test battery by Lewandowsky et al. (2010) implemented in MATLAB. Participants had to remember and recall letters in order of presentation. Letters were presented in set sizes of three to seven. Each set size occurred three times. As distractor task, before each letter was presented, a sentence was displayed that had to be classified as “correct” or “false” (e.g. “The earth is larger than the sun.”). The difficulty in this distractor task was kept low because this had improved the correspondence between the Sentence Span measure and a latent measure of working memory capacity in a previous study Lewandowsky et al. (2010).

### Stimuli for EEG Experiment

In the main EEG experiment, participants listened to natural speech in background noise that was made up of BT. The speech material was taken from a recording of a German audio book *(Die Glasglocke* by Sylvia Plath), recorded by a professional female speaker (F0 = 160.47 Hz, sd = 8.91 Hz).

First, the recording was manually split into segments that were coherent in content and had a length of about 45s. Second, these longer segments were split into three shorter segments of about 15 seconds (mean duration = 14.69 s, sd duration = 3.46 s). Special care was taken to ensure that all of these three shorter segments ended with a full stop.

**Table 2.**
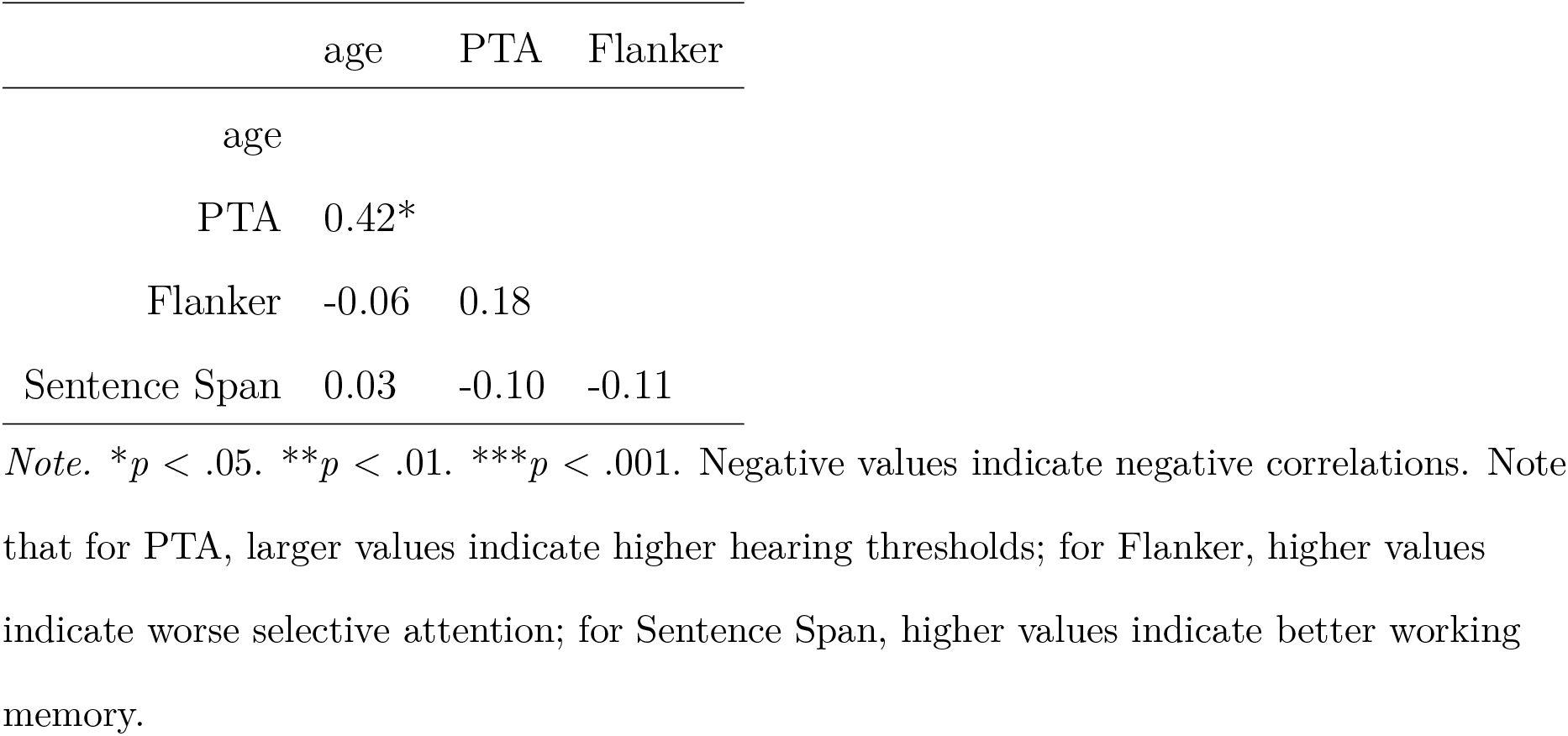
Correlation matrix of age and envelope tracking predictors

To create the speech-in-noise stimuli, we first trimmed all the silent periods in the longer sound segments and normalized them to 70 dB. Then, we mixed two or eight of these segments together to create background noise for the 2 BT and the 8 BT conditions. This mixture was then again normalized to 70 dB. After this, we manipulated the sound files to fade in over the first 1.5s. Afterwards, we mixed the background noise with the target speech segments at an SNR of either 0 or 2 so that the background noise would fade in after 2s of only the target speaker talking. This progressive addition of the background noise was implemented because target and noise speech signals were voiced by the same female speaker, and participants would otherwise not have been able to follow the target signal. Finally, this mixture was again normalized to 70 dB.

To create probe stimuli for a pattern-matching task, a short snippet of 0.3s was extracted from the last sixth of each sound file. This way, we ensured that the participants would continually need to track the target speaker to correctly categorize the probe snippet. We also ensured that the probe snippet would contain continuous speech and not a pause in the speech signal.

### Speech-in-Noise Tasks

Participants completed four experimental blocks. Each block contained 39 trials of a single condition. Trials length varied between 7.29 and 28.54s. Each sound segment was presented without background noise for 2s, after which the background noise faded in and ramped up until it reached its maximum sound level 3.5s after segment onset. An intelligibility task (IT) modeled after the pattern-matching task of (Liem et al., 2014) was implemented after the end of each trial. For the IT, a probe stimulus (the short sound snipped of 0.3s duration) was played, which was taken either from the sound segment or from the one of the BT. By means of a mouse click, participants stated whether the probe had been taken from the to-be-attended sound segment or not. If participants did not answer for 3s, the next trial began. Stimulus presentation was controlled via sound card (RME Babyface Pro, RME, Haimhausen, Germany) and stimuli were presented via a loudspeaker with linear frequency response (8030B Studio Monitor, Genelec, Iisalmi, Finland).

Each conglomerate of three IT trials was taken from one of the longer, coherent sound segments from the audio book. After these three IT trials, a comprehension task (CT) trial followed. In the CT, participants answered a four-alternative choice question about the content of the three previous trials by means of a keyboard button press. The comprehension task was untimed, so that participants would not feel rushed to answer the question.

### EEG Recording and Preprocessing

Participants sat in an EEG cabin in front of a computer screen. After a short instruction in which they were shown their EEG on a screen and could try out blinking, closing their eyes and grinding their teeth, they were asked to refrain from moving as much as possible. Then, a total of four minutes of resting-state EEG was recorded (2 minutes with eyes open, 2 minutes with eyes closed). In the eyes open condition, participants were asked to fixate a fixation cross on the computer screen.

The practice block contained three easy IT trials with 8 BT and an SNR of 5, followed by one CT question pertaining to the content of the three IT trials. In case the participant gave a wrong answer in the practice IT, that trial was repeated until the correct answer was given. After the practice session, participants were encouraged to attenuate or amplify the stimuli in order to ensure proper audibility. The original loudness of the stimuli was 70 dB SPL, and the range of attenuation/amplification across participants was −5 to 2 dB. Therefore, the final loudness ranged between 65 and 72 dB SPL. Critically, participants’ attenuation/amplification was correlated with their PTA (*r* = −0.56, *p* = 0.004), with participants with higher PTA aiming for louder stimuli. After stimulus loudness adjustment, participants completed four experimental blocks, one for each condition, each of which took about 15 minutes. Block order was counterbalanced between participants.

Participants’ EEG was recorded continuously from 128 Ag/AgCl electrodes (BioSemi ActiveTwo, Amsterdam, The Netherlands) with a ActiveTwo AD-box amplifier system (BioSemi ActiveTwo, Amsterdam, The Netherlands) and was digitized at a sampling rate of 512 Hz. The data were online band-pass filtered between 0.1–100 Hz and impedances were reduced below 25 kOhm. Data were analyzed in MATLAB Release 2016b (The MathWorks, Inc., Natick, Massachusetts, United States) using the FieldTrip Toolbox (Oostenveld et al., 2011). For pre-processing, data were re-referenced to Cz and then band-pass filtered between 0.1 and 100 Hz with a non-causal zero-phase two-pass 4th order Butterworth IIR filter with −12 dB half-amplitude cutoff. A non-causal zero-phase two-pass 4th order Butterworth IIR band-stop filter with −12 dB half-amplitude cutoff was applied between 48 and 52 Hz in order to eliminate artifacts resulting from electric interference. Data were visually screened for noisy channels, which were then removed. After that, the continuous EEG was segmented into trials starting 2s before sentence onset and lasting until the end of the sentence (mean trial duration = 16.69 s, sd = 3.46 s). Trials containing gross artifacts were removed. After that, data were re-referenced to an average reference and an independent component analysis (ICA) (Jung et al., 2000) was applied. For the ICA, data were high-pass filtered at 1 Hz in order to improve stationarity of the components. After the removal of artefactual components, the remaining components were back-projected to the original, 0.1-Hz-filtered data. Finally, noisy channels were interpolated using spline interpolation (Perrin et al., 1987).

### Temporal Response Function

We used the mTRF toolbox (Crosse et al., 2016) and followed the recommendations in the documentation in order to fit Temporal Reponse Functions (TFRs). We extracted the envelopes of the target speech signals with the *mTFRenvelope* function, downsampled them to 128 Hz and z-scored them. We also downsampled the EEG of each trial to 128 Hz and z-scored it. Additionally, the first 3.5s of each EEG and envelope were removed because the background noise was either missing or ramping up during that time. Conveniently, this step also removed any activity in relation to early cortical evoked potentials. We identified the optimal ridge parameter via the *mTRFcrossval* function. Then, a model for each condition, each channel, and each participant was trained via the *mTRFtrain* function, again with the z-scored EEGs and envelopes. Each model was calculated over time lags between envelope and EEG between −150 and 450 ms, after which values for lags from −150 to −100 and from 400 to 450 ms were removed because of regression artifacts at the extremes of the models. For each trials, we additionally fitted TRFs between that trial’s EEG and the envelope of the following trial (the EEG of each last trial per condition was paired with the envelope of the first trial). This was done in order to obtain baseline TRFs, which would be contrasted with the actual TRFs as a measurement for TRF quality.

### Statistical Analysis

Statistical tests were conducted in R, Version 3.6.2 (Team, 2018) and FieldTrip, Version 20190419 (Oostenveld et al., 2011). The *p*-values for estimates in linear mixed-effects models (LMEM) were derived via the Satterthwaite method implemented in the R package *lmerTest* (Kuznetsova et al., 2017).

All TRF analyses took place at lags between −100 and 400 ms. All statistical comparisons between conditions were conducted using cluster-based permutation tests implemented in Fieldtrip (Maris and Oostenveld, 2007). Dependent-samples t-tests were conducted for each sample between the TFRs of the two conditions to be compared. All samples whose t-value exceeded a p-value of 0.05 were clustered on the basis of temporal adjacency, with a cluster containing at least three neighboring channels. This was done separately for samples with negative and positive t-values. The t-values of each cluster were summed and the maximum absolute value of the cluster-level statistics was taken. This maximum absolute value was then compared to a permutation distribution, which was obtained by randomly assigning the TFRs to the compared conditions, calculating the test statistic on this random set of trials and repeating this procedure 1000 times. The critical *α* level for comparing the test statistic to the permutation distribution was set to 0.025 for a two-sided t-test Maris and Oostenveld (2007). All cluster-based tests reported followed this procedure.

## Results

### Behavioral Results

Please see Table 3 for an overview of performance in the behavioral tasks. To quantify performance in the IT, we calculated d’. We then fitted a LMEM predicting d’ from SNR and nBT and their interaction, with a random intercept for participant (random slopes models did not converge). There was no significant effect of the condition factors nor their interaction. Because d’ was rather low for all conditions, due to a high rate of false alarms, we fit the same model to the hit rate and the false alarm rate separately. SNR significantly predicted hit rate, *b* = 0.13, *t*(66) = 4.49, *p* < .001, with SNR 2 resulting in a higher hit rate than SNR 0. However, SNR also significantly predicted the false alarm rate, *b* = 0.10, *t*(66) = 3.06, *p* = .003, again with SNR 2 resulting in a higher false alarm rate. There was likely no significant effect of SNR on d’ because these two effects cancelled each other out.

**Table 3.**
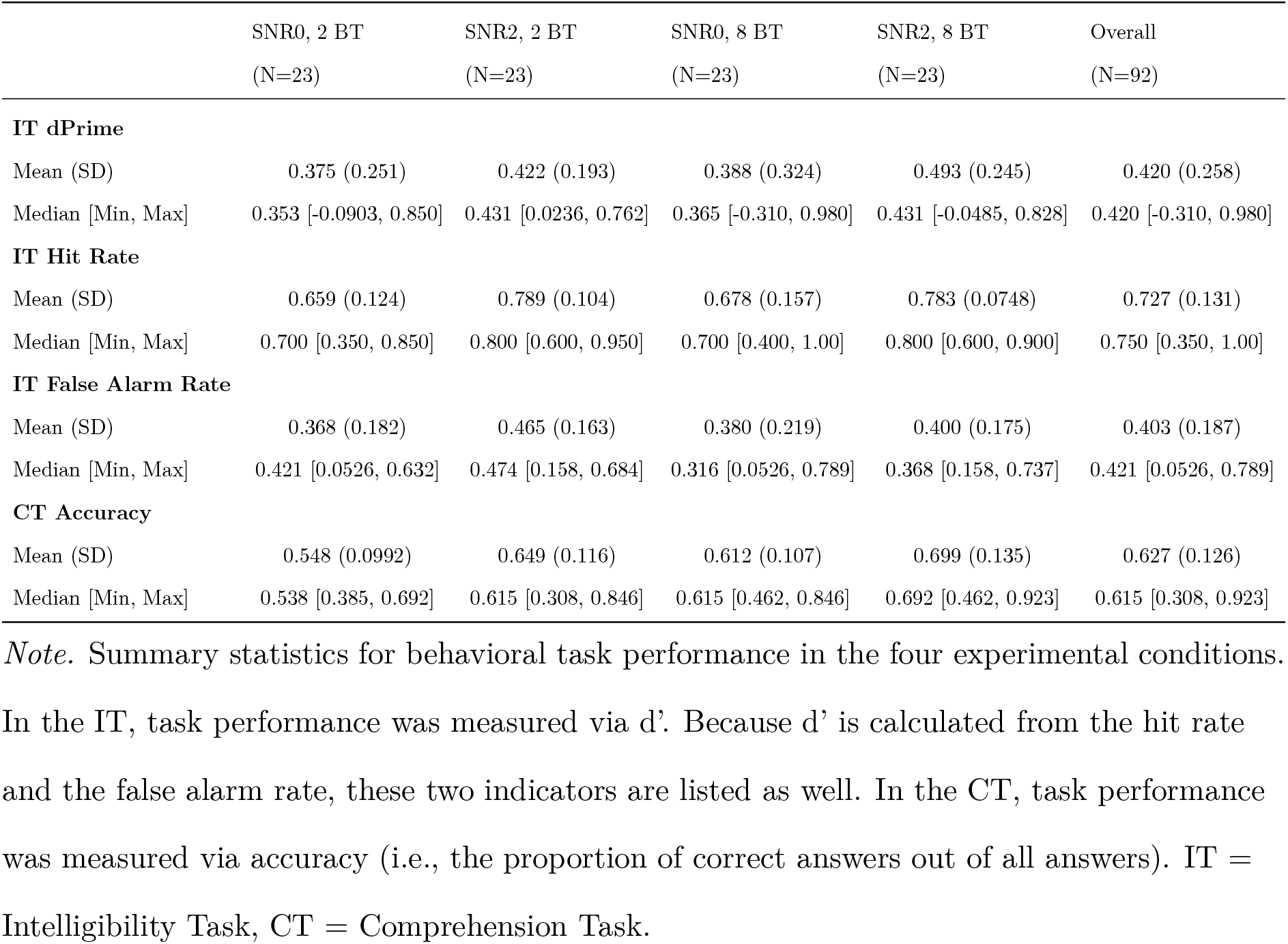
Summary Statistics of Task Performance

|  | SNR0, 2 BT<br>(N=23) | SNR2, 2 BT<br>(N=23) | SNR0, 8 BT<br>(N=23) | SNR2, 8 BT<br>(N=23) | Overall<br>(N=92) |
| --- | --- | --- | --- | --- | --- |
| **IT dPrime** |  |  |  |  |  |
| Mean (SD) | 0.375 (0.251) | 0.422 (0.193) | 0.388 (0.324) | 0.493 (0.245) | 0.420 (0.258) |
| Median [Min, Max] | 0.353 [-0.0903, 0.850] | 0.431 [0.0236, 0.762] | 0.365 [-0.310, 0.980] | 0.431 [-0.0485, 0.828] | 0.420 [-0.310, 0.980] |
| **IT Hit Rate** |  |  |  |  |  |
| Mean (SD) | 0.659 (0.124) | 0.789 (0.104) | 0.678 (0.157) | 0.783 (0.0748) | 0.727 (0.131) |
| Median [Min, Max] | 0.700 [0.350, 0.850] | 0.800 [0.600, 0.950] | 0.700 [0.400, 1.00] | 0.800 [0.600, 0.900] | 0.750 [0.350, 1.00] |
| **IT False Alarm Rate** |  |  |  |  |  |
| Mean (SD) | 0.368 (0.182) | 0.465 (0.163) | 0.380 (0.219) | 0.400 (0.175) | 0.403 (0.187) |
| Median [Min, Max] | 0.421 [0.0526, 0.632] | 0.474 [0.158, 0.684] | 0.316 [0.0526, 0.789] | 0.368 [0.158, 0.737] | 0.421 [0.0526, 0.789] |
| **CT Accuracy** |  |  |  |  |  |
| Mean (SD) | 0.548 (0.0992) | 0.649 (0.116) | 0.612 (0.107) | 0.699 (0.135) | 0.627 (0.126) |
| Median [Min, Max] | 0.538 [0.385, 0.692] | 0.615 [0.308, 0.846] | 0.615 [0.462, 0.846] | 0.692 [0.462, 0.923] | 0.615 [0.308, 0.923] |

*Note.* Summary statistics for behavioral task performance in the four experimental conditions.
In the IT, task performance was measured via $d'$ . Because $d'$ is calculated from the hit rate and the false alarm rate, these two indicators are listed as well. In the CT, task performance was measured via accuracy (i.e., the proportion of correct answers out of all answers). IT = Intelligibility Task, CT = Comprehension Task.

We also fitted a LMEM to the accuracy of the CT with SNR, number of talkers, and their interaction as predictors and a random intercept of participant. SNR significantly predicted accuracy in the CT, *b* = 0.10, *t*(66) = 3.19, *p* = .002, with higher accuracy for the SNR 2 conditions than for the SNR 0 conditions. Also, nBT significantly predicted accuracy in the CT, *b* = 0.06, *t*(69) = 2.02, *p* = .048, with a higher accuracy for the conditions with 8 BT than for the conditions with 2 BT. Therefore, both SNR and nBT influenced performance in the CT, but there was no evidence for an interaction effect of the two.

### TRF Results

Figure 1 shows the grand average TRFs for the four conditions and the grand average baseline TRF. Viewing the TRFs, in contrast to earlier MEG TRF estimations, we not only observed peaks in the TRF at around 50 and 100 ms, but also a third, prolonged peak, similar to the Pd in Power et al. (2012) and the *P*2*_c_rosscorr* in Petersen et al. (2017). In reference to the approximate timing of their maximum deflection, we will refer to them as TRF_50_, TRF_100_, and TRF_300_.

**Figure 1.**
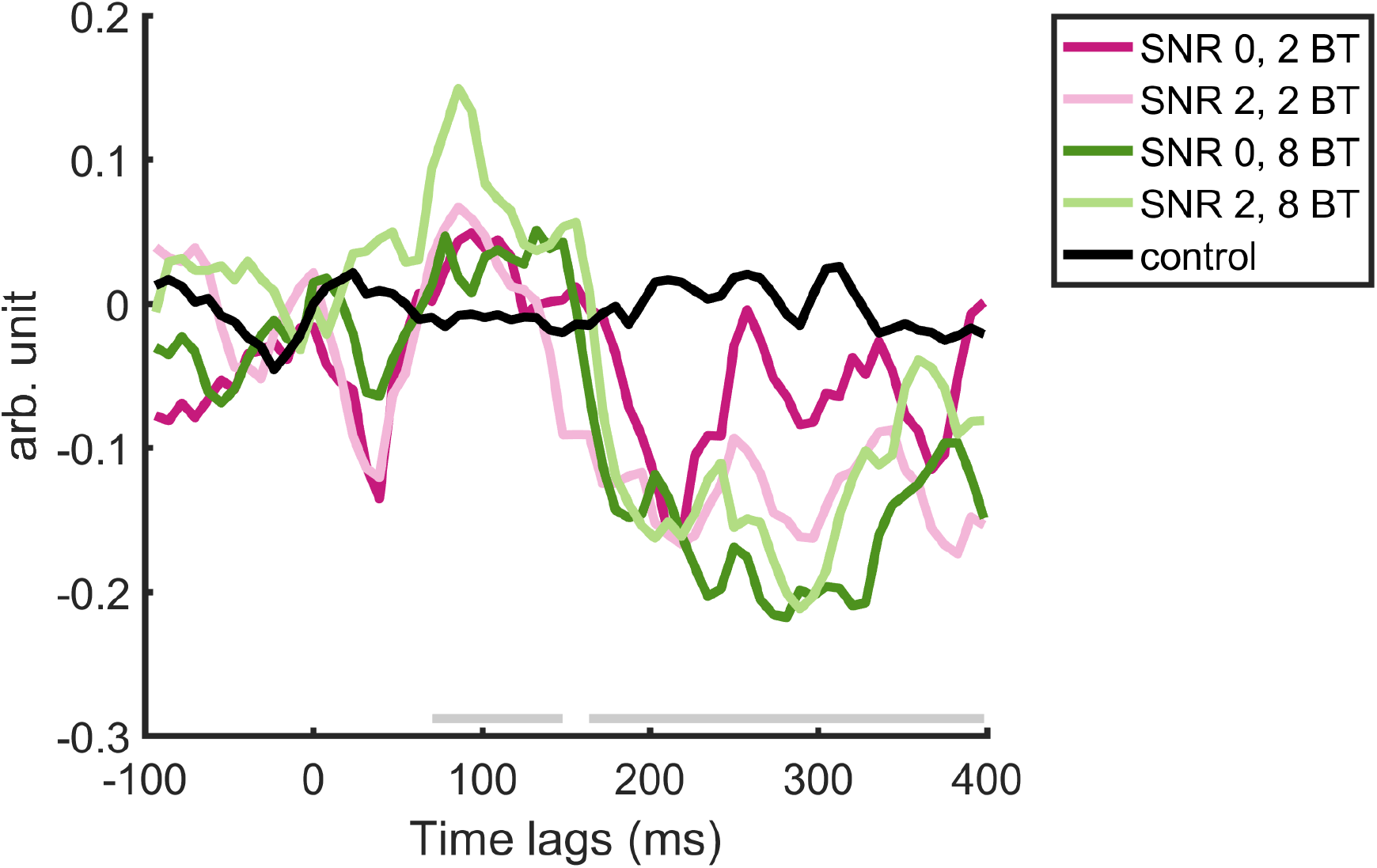
Grand average TFR traces of the four experimental conditions and the baseline TRF, averaged across postero-occipital midline electrodes (A21, A22, A23, A24). Time lags at which the average of the actual TRFs significantly differed from the baseline TRF (*p* < 0.05) are denoted with the gray bar slightly above the x axis.

Using cluster-based permutation tests, we first compared all TRFs to the baseline TRFs. All TRFs when compared to the baseline separately differed significantly in a first time window starting from ~ 70ms until ~ 150 ms and in a second, longer time window, starting from ~ 170 ms and lasting until the end of our time window. These differences were significant at almost all electrodes. We therefore concluded that TRF estimation had been successful in capturing brain activity related to the speech envelope. There was a small window between lags of around 150 and 170 ms where no significant difference was found. We suspect that this is due to the reversing of the polarity during this time window, which is bound to cross the zero line, around which the control TRFs hovered. Because sound onsets, which would have elicited auditory ERPs, had been removed from the EEG with which the TRFs were calculated, our TRFs represent brain activity related to the speech envelope (linearly; Crosse et al., 2016).

#### TRFs as a Function of SNR and nBT

Next, we compared the TRFs of conditions with SNR 0 to the TRFs of conditions with SNR 2. There was no significant difference at any time lag between the two conditions.

Then, we compared the TRFs of the 2 BT conditions to the TRFs of the 8 BT conditions. There were two time lag windows at which the TRFs were significantly different; one early time window at lags of around 10 to 60 ms, reflecting TRF_50_ and one later time window at lags of around 210 to 330 ms, reflecting TRF_300_ (see Figure 2). Inspection of the topography at significantly different time lags (see also Figure 2) revealed that these differences came about because of one positive and one negative cluster at each time lag. At each of the significantly different time windows, one negative and one positive cluster complemented each other, with one being located at left anterior and medial anterior electrodes and the other at postero-occipital electrodes. Figure 2 shows the TRFs and topoplots of the clusters. Note that the terms “positive” and “negative” refer to the difference in t-values between the conditions, and not to the polarity of the peaks associated with them. In fact, the positive clusters reflected negative peaks and the negative clusters reflected positive peaks in the TRFs. Because of that, more negative values in the positive clusters reflect larger amplitudes in the peaks associated with it, as do more positive values in the negative clusters.

**Figure 2.**
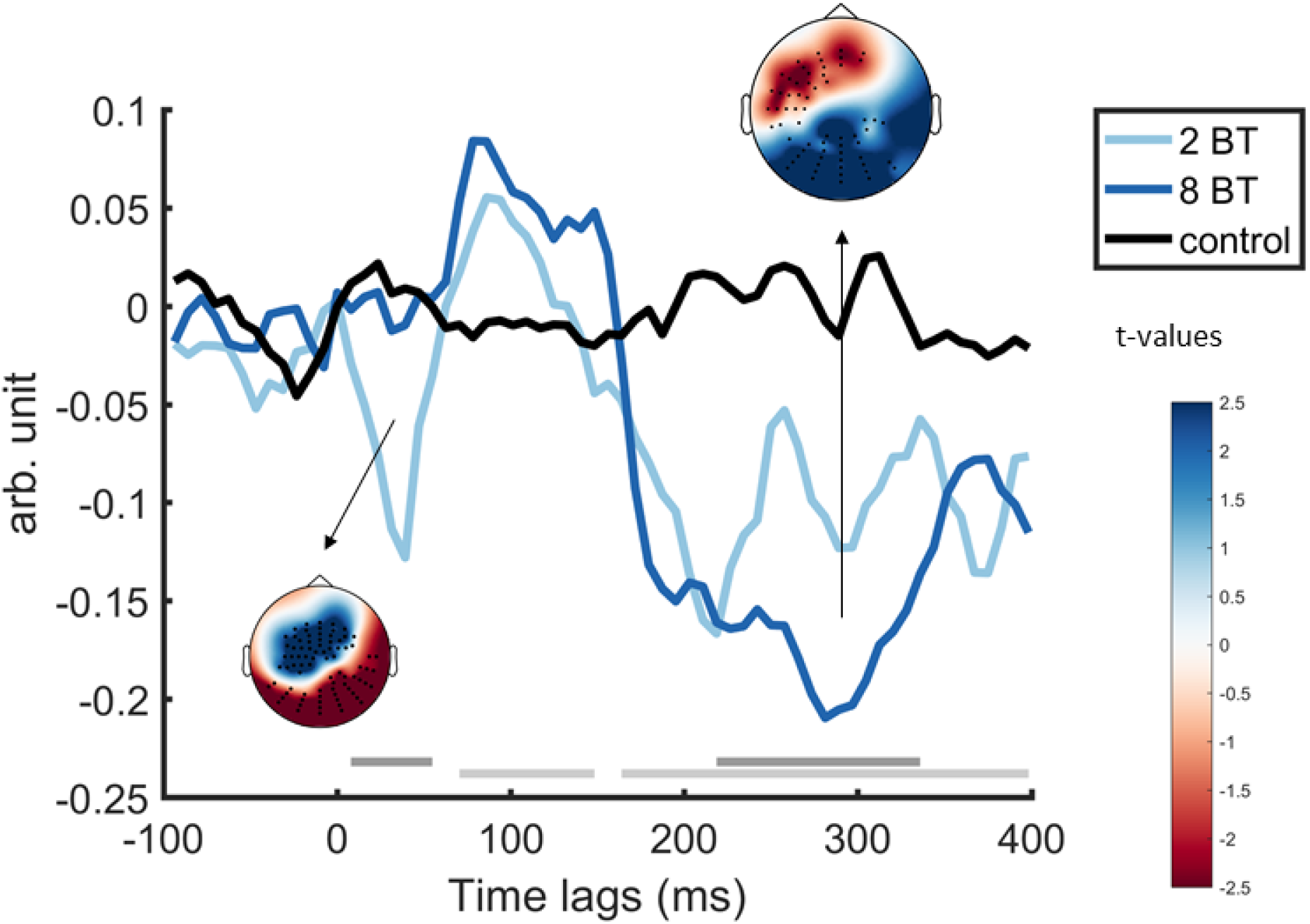
Grand average TFR traces of the 2 BT conditions irrespective of SNR, the 8 BT conditions irrespective of SNR, and the baseline TRF, averaged across postero-occipital midline electrodes (A21, A22, A23, A24). Time lags at which the average of the actual TRFs differed from the baseline TRF are denoted with the light gray bar slightly above the x axis. Time lags at which the 2 BT TRFs differed from the 8 BT TRFs are denoted with the darker gray bar. Additionally, the plot shows the topographies of t-values at 31.3 ms and at 281 ms time lags.

As a next step, we tested the interaction effect of SNR and nBT. For that, we performed cluster-based comparisons between the raw effect of SNR (difference in SNR TFRs) and the raw effect of nBT (difference in BT TRFs). There was no significant difference at any time lag in the interaction of the two conditions.

### TRF cluster peak amplitudes as a function of SNR, nBT, hearing and cognition

To test whether there was an association between TRF peak amplitudes and inter-individual variables of hearing and cognition, we exported average values for the two positive and the two negative clusters that reached significance in the 2 vs. 8 BT comparison. While reducing these multidimensional data to a single value means losing temporal and spatial information, it makes the fitting of more complex models possible.

#### Cooccurrence of Clusters

First, we asserted whether the presumed association between the simultaneously occurring negative and positive clusters could be confirmed with the averaged values. To this end, we ran Person correlations between exported values of the positive and negative clusters of TRF_50_ and the positive and negative clusters of TRF_300_. The amplitudes in the TRF_50_ clusters were highly correlated (*r*(90) = −0.84, *p* < 0.001), as were the amplitudes in the TRF_300_ clusters (*r*(90) = −0.78, *p* > .001). These strong correlations support the view that positive and negative clusters occurring at the same time represent the same or at least related processes.

#### TRF Amplitude as a Function of SNR and nBT

Second, we aimed to replicate the result of the FieldTrip analysis within R by fitting LMEMs with SNR and nBT and their interaction effect as fixed effects and a random intercept per participant (a random slopes model did not converge) to each of the four average cluster values. All four of the models contained a significant main effect of nBT (positive cluster of TRF_50_: *b* = −0.08, *t*(66) = −3.52, *p* < .001; negative cluster of TRF_50_: *b* = 0.08, *t*(66) = 3.01, *p* = .004; positive cluster of TRF_300_: *b* = −0.15, *t*(66) = −5.45, *p* < .001; negative cluster of TRF_300_: *b* = 0.12, *t*(66) = 5.27, *p* < .001), with the amplitude being higher in the 8 BT conditions than in the 2 BT conditions. Additionally, for the TRF_300_ positive cluster, there was a trend for a main effect of SNR, *b* = −0.05, *t*(66) = −2, *p* = .05, with a higher amplitude in the SNR 2 conditions than in the SNR 0 conditions.

Furthermore, for the TRF_300_ positive cluster, there was a significant interaction effect of SNR and nBT, *b* = 0.08, *t*(66) = 2.13, *p* < .04. Although amplitude was always higher in the 8 BT conditions than in the 2 BT conditions, the difference between the two was stronger when the SNR was 0 than when the SNR was 2.

#### TRF Amplitude as a Function of SNR, nBT, and Participant-Level Variables

In a third step, we updated the LMEMs with participant-level variables. Specifically, we each added PTA, Sentence Span for working memory, and Flanker for selective attention as z-scored predictors to the models separately. We then performed likelihood ratio tests between the models with and without participant-level predictor. Neither the inclusion of PTA nor of working memory provided a significantly better fit to the data. The inclusion of selective attention provided a better fit to the data for models of the TRF_300_ positive cluster values, *χ*^2^(4) = 11.90, *p* = .02, and negative cluster values, *χ*^2^(4) = 10.24, *p* = .04. The inclusion of selective attention did not result in an additional significant effect in the model for the negative cluster values. For the positive cluster values, we found a three-way interaction effect between SNR, nBT, and selective attention. Model parameters are reported in Table 4 and the effects are visualized in Figure 3. TRF_300_ amplitude was always larger in the 8 BT conditions than in the 2 BT conditions, but this difference was larger in the SNR 0 conditions than in the SNR 2 conditions. There was also a significant interaction effect between nBT and selective attention, with participants with better selective attention (lower z-scores) having a stronger increase in amplitude between 2 BT and 8 BT. Regarding the three-way interaction, in the SNR 0 conditions, better selective attention (lower z-scores) led to a steeper decrease in TRF amplitude than worse selective attention. However, in the SNR 2 conditions, the increase in TRF_300_ amplitude between the 2 BT and the 8 BT conditions was steeper for participants with worse selective attention.

**Figure 3.**
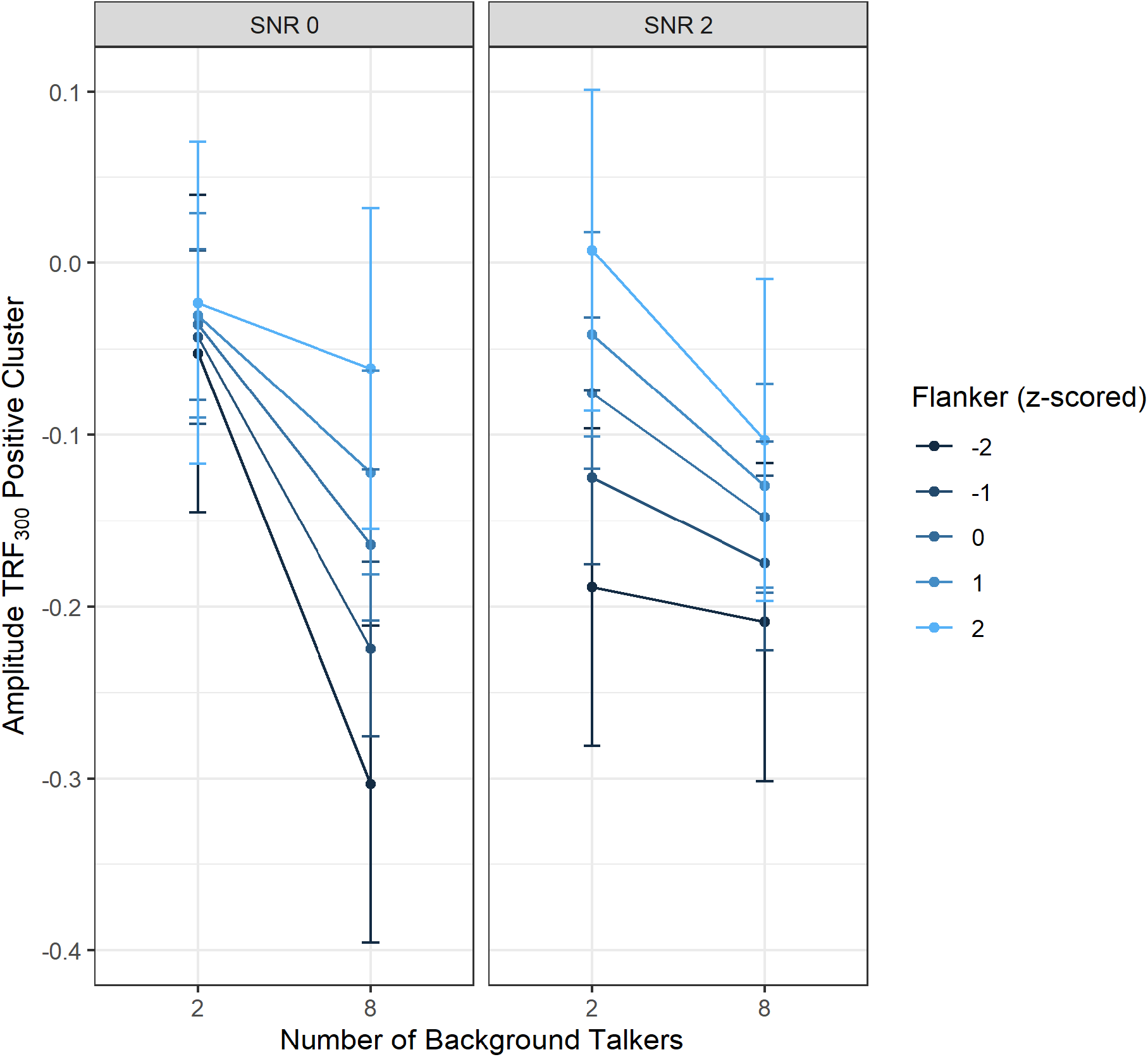
Three-way interaction effects plot of SNR, nBT, and selective attention. Bars indicate 95% confidence intervals.

**Table 4.**
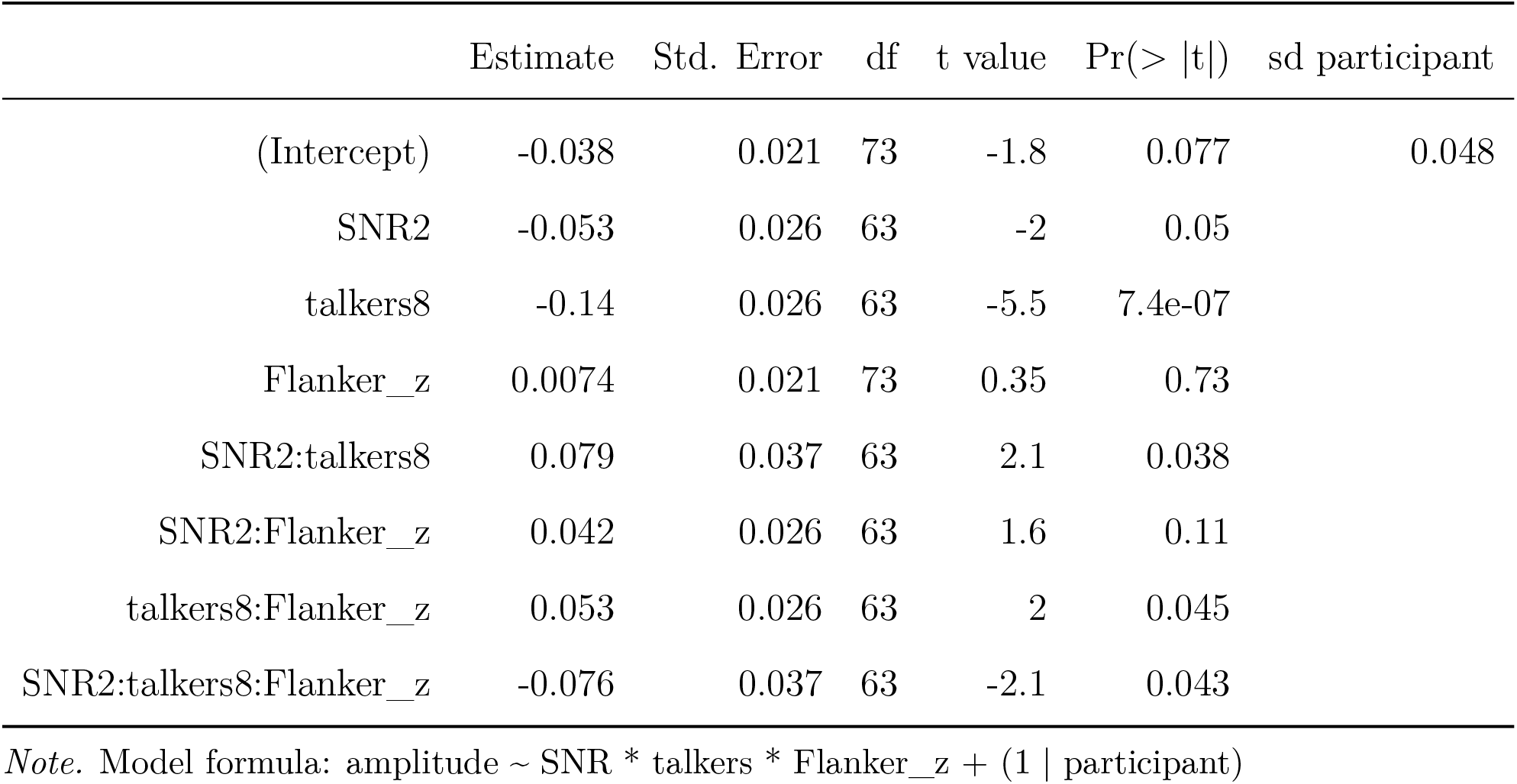
Parameters from model predicting TRF_300_ positive cluster amplitude from SNR, nBT, and selective attention.

|  | Estimate | Std. Error | df | t value | Pr(> \|t\|) | sd participant |
| --- | --- | --- | --- | --- | --- | --- |
| (Intercept) | -0.038 | 0.021 | 73 | -1.8 | 0.077 | 0.048 |
| SNR2 | -0.053 | 0.026 | 63 | -2 | 0.05 |  |
| talkers8 | -0.14 | 0.026 | 63 | -5.5 | 7.4e-07 |  |
| Flanker_z | 0.0074 | 0.021 | 73 | 0.35 | 0.73 |  |
| SNR2:talkers8 | 0.079 | 0.037 | 63 | 2.1 | 0.038 |  |
| SNR2:Flanker_z | 0.042 | 0.026 | 63 | 1.6 | 0.11 |  |
| talkers8:Flanker_z | 0.053 | 0.026 | 63 | 2 | 0.045 |  |
| SNR2:talkers8:Flanker_z | -0.076 | 0.037 | 63 | -2.1 | 0.043 |  |

*Note.* Model formula: amplitude ~ SNR \* talkers \* Flanker\_z + (1 | participant)

### Task performance as a Function of SNR, nBT, and Cluster Amplitudes

We further tested whether the inclusion of cluster amplitude improved the models for performance in the IT and the CT task. To this end, we updated the previous models on hit rate and false alarm rate in the IT and accuracy in the CT to include each of the positive and negative TRF_50_ and TRF_300_ clusters in separate models with full main and interaction effects with SNR and nBT. In total, 12 models were fitted. Again, we first tested whether this addition would provide a better fit to the data with likelihood ratio tests.

#### IT Task Performance as a Function of SNR, nBT, and Cluster Amplitudes

For the IT task, the inclusion of the TRF_300_ positive cluster into the model for the hit rate provided a significantly better fit to the data than the basic model with just SNR and nBT, *χ*^2^(4) = 11.44, *p* = .02. Also, the inclusion of the TRF_300_ negative cluster provided a significantly better fit, *χ^2^*(4) = 11.38, *p* = .02. However, no additional significant effects were found in the models. Inclusion of the TRF_50_ or TRF_300_ cluster values did not improve model fit for the false alarm rate of the IT.

#### CT Task Performance as a Function of SNR, nBT, and Cluster Amplitudes

For accuracy in the CT, the inclusion of the TRF_300_ positive cluster into the model for the hit rate provided a significantly better fit to the data than the basic model, *χ*^2^(4) = 11.39, *p* = .02. However, this did not result in an additional significant effect in the model. The inclusion of the TRF_300_ negative cluster into the model for accuracy in the CT provided a significantly better fit, *χ*^2^(4) = 15.24, *p* = .004. Amplitude of the TRF_300_ negative cluster significantly predicted accuracy in the CT, *b* = 0.13, *t*(83.17) = 2.59, *p* = .01, with a larger amplitude (more positive values) resulting in better accuracy. Additionally, there was a significant interaction effect between SNR and negative cluster amplitudes, *b* = −0.12, *t*(81.94) = −2.03, *p* = 0.046. This interaction effect is visualized in Figure 4. Amplitude of the TRF_300_ negative cluster predicts accuracy in the SNR 0 conditions, with a higher amplitude being related to higher accuracy, but not in the SNR 2 conditions.

**Figure 4.**
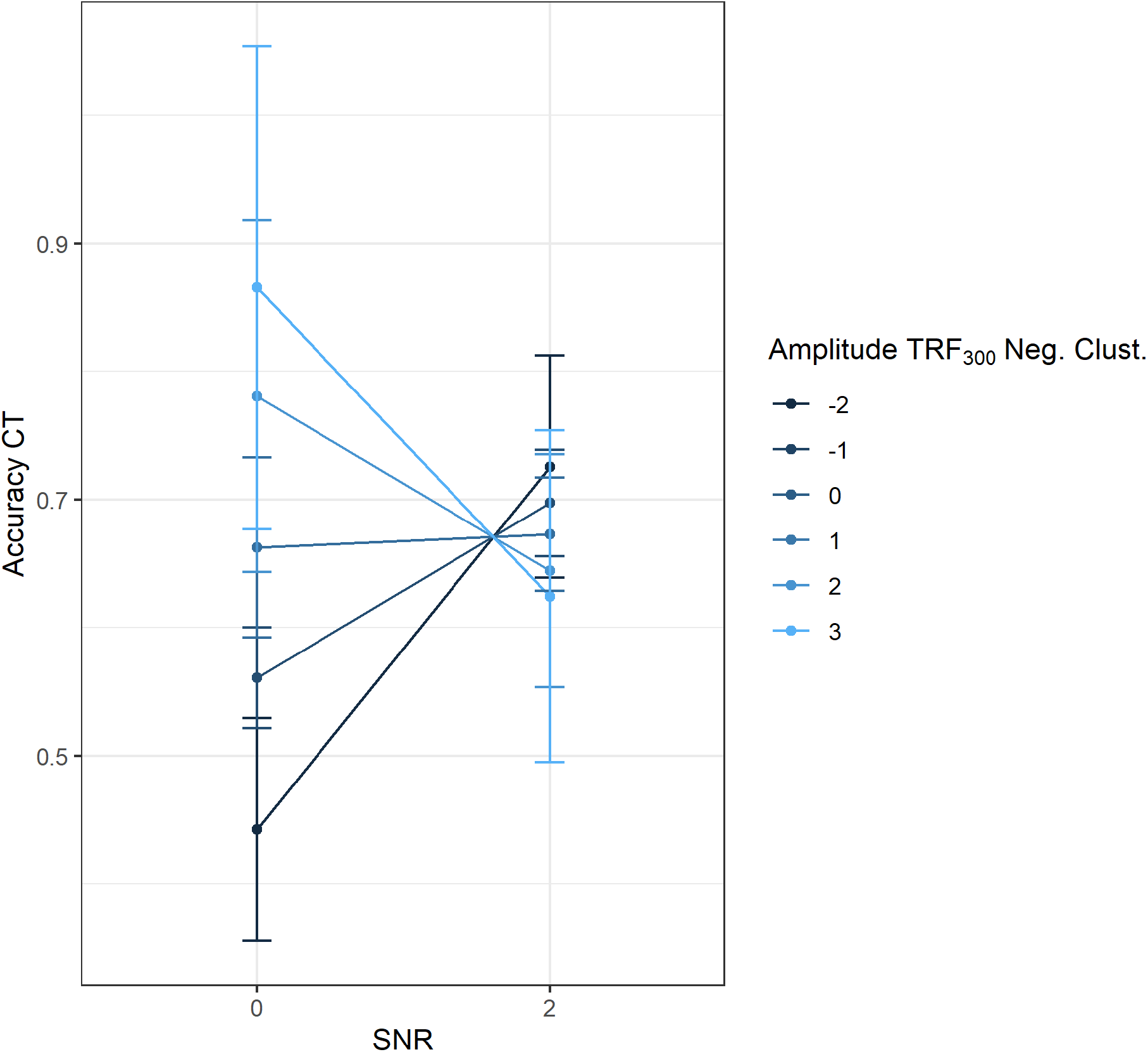
Effects plot of the interaction between SNR and amplitude of the TRF_300_ negative cluster. Bars indicate 95% confidence intervals.

## Discussion

The study aimed to investigate neural envelope tracking in multi-talker babble noise in healthy older adults, how it is affected by varying degrees of EM and IM, and how peripheral hearing and cognition modulate envelope tracking in these situations.

### Behavioral Task Performance

At a behavioral level, we mainly observed effects of EM and IM in the CT, with higher SNR and higher number of talkers resulting in a higher accuracy. In the IT, the higher hit rate in the higher SNR conditions was cancelled out by the higher false alarm rate in the same conditions. It is unclear why a higher SNR should result in a higher false alarm rate. Intuitively, one would expect the opposite. A similar effect was found in the study by Brungart (2001), who observed a u-shaped effect of SNR on call sign identifications when the background noise consisted of the same talker as the target talker. In their study, performance was lowest when the SNR was 0, and, as expected, increased when the SNR was above 0. Surprisingly, the performance also improved when the SNR decreased below 0. Brungart (2001) explained this phenomenon with the increased dissimilarity between speakers of different intensity levels, which leads to a reduction in IM and outweighs the additional EM that results from the decreased SNR. However, in our study, false alarm rate was not highest at SNR 0, but at SNR 2, and, therefore, cannot be explained with an inadvertent increase in IM due to similarity. The pattern rather suggests that the SNR 2 conditions fostered an answer bias and that answering the pattern-matching prompt with ‘yes’ occurred more often in the SNR 2 than in the SNR 0 conditions. Nevertheless, our behavioral results can be viewed within the effortfulness hypothesis (e.g. McCoy et al., 2005), which sees decrements in performance not necessarily at early stages (here following the target speaker in the IT), but when taxed with additional working memory load (answering content questions in the CT).

### Envelope Tracking in Energetic and Informational Masking

The main research question of the study was how EM and IM independently and jointly influence envelope tracking in older adults. With regard to EM, we found no effect of SNR on TRFs. Our results concur with those of a study that did not find an effect of SNR on TRFs as well (Ding and Simon, 2012). Although we had expected an early effect of SNR, as had been found in other studies, these studies either presented stimuli with very salient SNR differences (Ding and Simon, 2013) or presented SNR differences specific to participants’ SRTs (Petersen et al., 2017). Possibly, with greater SNR differences, this effect might have emerged in our sample as well. Unfortunately, our SNR range was limited as pilot testing revealed that a higher SNR would have resulted in ceiling performance in the behavioral tasks. Indeed, when reducing the data to an average value across the time of the TRF_300_ peak, there was an almost significant trend (*p* = 0.05) for a main effect of SNR, although in a later time window than expected. Also, as more variables are introduced, SNR differences do seem to be meaningful, as will be explored in the corresponding sections of the discussion.

There was a significant difference in the TRFs to the 2 vs. 8 BT conditions. Both the TRF_50_ and the sustained later peak, the TRF_300_, exhibited a larger amplitude in the 2 BT conditions than in the 8 BT conditions. In an attended/competing talker paradigm, Power et al. (2012) conducted a TRF/AESPA analysis and found an attention-related component at around a lag of about 220 ms, the Pd, which was present for the speech envelope of an attended talker, but not for the envelope of the competing talker, and which they related to a semantic filtering process. Another study (Niemczak and Vander Werff, 2019) investigated how the P1-N1-P2 response was affected by EM and IM. In this study, IM was also manipulated by varying the nBT between 2 BT and 8 BT. They found a reduced P2 amplitude in the 2 BT condition compared to the 8 BT condition. Lalor et al. (2009) already assumed that the Pd (corresponding to our TRF_300_ cluster) might be related to the P2. Our results add to this body of research in supporting that a deflection in this time window might indeed reflect semantic filtering, based on the fact that our IM stimuli were constructed to reflect different amounts of perceivable semantic content.

The finding of the TRF_50_ cluster is especially interesting, because we did not expect such an early modulation of target signal envelope tracking between these two experimental conditions. By reducing glimpsing opportunities through elimination of silent periods in the background talkers’ speech signals (see Materials and Methods section) and by creating background noise taken from the same speaker as the target speaker, we explicitly targeted lexical interference as the main component of interest of IM. Nevertheless, it is possible that stronger envelope tracking (as reflected in the larger TRF_50_) occurred in the 2 BT conditions because of remaining glimpsing opportunities. On another note, it might be useful to complement the analysis with an account of the auditory ERP component corresponding to the TRF_50_ with regard to temporal occurrence, the P50. The P50 is involved in sensory gating (Joos et al., 2014) and it has been suggested that top-down modulation of sensory input is already present during such an early time window (Kurthen et al., 2007). Possibly, object formation (Shinn-Cunningham, 2008) was more prominent in the 8 BT conditions because more auditory objects were present (there were nine instead of three speakers to encode), which in turn attenuated the response to each single speaker, including the target speaker.

The finding of the significant difference in the TRF_300_ cluster was expected because it occurred in a time window where an involvement of cognitive ability in stimulus processing had already been demonstrated (van Dinteren et al., 2014; Giroud et al., 2016; Snyder et al., 2006; Picton and Hillyard, 1974). When visually examining the two TRF traces in Figure 2, it seems that the TRF_300_ is not only of reduced amplitude but also of shorter duration in the 2 BT conditions than in the 8 BT conditions. Given that in the study by Power et al. (2012), an AESPA peak at a similar latency signalled attention, this could reflect reduced attention directed at the target speaker in the 2 BT conditions. This interpretation is corroborated by the significant interaction between nBT and Flanker performance in the model that predicted the amplitude of the positive TRF_300_ cluster.

Using cluster-based permutation tests, we found no interaction effect of SNR and nBT and thus no evidence that EM and IM add up in their effects on speech envelope tracking at any point.

### TRF Peak Amplitudes as a Function of Experimental Condition, Hearing, and Cognition

Because envelope tracking is assumed to take on a functional role for speech understanding (Riecke et al., 2018; Wilsch et al., 2018; Zoefel et al., 2018), it is important to investigate how envelope tracking is affected by inter-individual participant characteristics. Identifying inter-individual variables that influence envelope tracking might also explain differences in speech understanding.

In our study, we investigated how hearing thresholds, working memory, and selective attention influence speech envelope tracking. Neither hearing thresholds nor working memory were shown to predict envelope tracking. This was surprising given that hearing thresholds have been an important predictor for TRF peak amplitude in the study of Petersen et al. (2017). However, in their study, it predicted the amplitude of the *N*1*_crosscorr_*, which did not show up as a significantly different peak in our analysis and was therefore not subjected to such an analysis in our study. Additionally, we accounted for inter-individual differences in hearing thresholds by allowing participants to alter the sound level of the stimuli. Possibly, hearing thresholds play a significant role in early, perceptual stages of speech processing, but not as much in later, cognitive stages. Consequently, counteracting peripheral hearing loss by means of a hearing aid might aid the early, perceptual processing of a speech signal, but might not be as effective in supporting the later, cognitive processing of speech in background talker noise (Bertoli et al., 2009).

As for working memory, we would have expected it to predict TRF amplitude at a later point in time because of theoretical reasons. Working memory is by far the most commonly found predictor for speech-in-noise processing, and in the study by Decruy et al. (2019), working memory was positively related to envelope tracking in the presence of a competing talker, but this information stems from a significant interaction effect between background noise type and working memory, and not from a main effect of working memory itself. Also, they used a backward modeling approach which allowed for a quantification of the reconstruction of the envelope, but not for its time course. Our approach of forward modeling allows for sample-to-sample comparison of conditions, and therefore enabled us to integrate previous knowledge of the involvement of cognition over the time course of speech processing into the investigation of EM and IM influences and thereby to identify two time windows (TRF_50_ and TRF_300_) during which differences in IM resulted in different amounts of envelope tracking. However, even with such temporal precision, there was no effect of working memory in our study. Our SNRs ranged from 0 to 2 dB, while in the study of Decruy et al. (2019), they ranged between 3 and −6 dB. Effects of working memory on envelope tracking between our conditions might have emerged with a larger difference in SNR.

The inclusion of selective attention as a variable provided a better fit to the data than just SNR and BT, but only for the TRF_300_ clusters. The three-way interaction between SNR, nBT, and selective attention illustrates the nontriviality of combining EM and IM. The inclusion of selective attention also revealed an interaction between SNR and nBT, which indicated that while cluster amplitude was always larger in the 8 BT conditions than in the 2 BT conditions, this difference was larger in the SNR 0 conditions than in the SNR 2 conditions. Therefore, stronger EM led to a greater difference in envelope tracking in conditions varying in IM. There was also a significant interaction effect between nBT and selective attention, with participants with better selective attention having a stronger increase in TRF_300_ amplitude between the two conditions. The three-way interaction revealed that this pattern was true only for the SNR 0 conditions, and that the increase in TRF_300_ amplitude between the two conditions was actually stronger for participants with worse selective attention in the SNR 2 conditions. These results can be interpreted as that the release from EM in the SNR 2 conditions played to the strengths of participants with lower selective attention ability, who, in the easier SNR conditions, could stronger differentiate between high and low IM. Contrary to our results, the study by Presacco et al. (2016a) found that selective attention was negatively related to envelope tracking in older adults. However, they measured envelope tracking by means of envelope reconstruction fidelity. With our forward-modeling approach, we could tap into envelope tracking during different time windows. Indeed, selective attention was only a relevant predictor in the later, TRF_300_ time window and not during the earlier TRF_50_ time window.

### Behavioral Relevance of Envelope Tracking

Finally, we were interested in whether envelope tracking served a functional role in speech understanding. We found envelope tracking in the later TRF_300_ time window to positively predict performance in the CT task, which measured how well participants could memorize the content of the target speaker’s speech signal. Additionally, the significant interaction effect between envelope tracking and SNR revealed that envelope tracking was even more positively related to CT task performance in the more difficult SNR 0 condition. Therefore, stronger envelope tracking seems especially helpful in a more difficult listening situation in terms of EM.

Additionally, relative to young adults, older adults on average exhibit stronger envelope tracking and it has been debated whether this stronger envelope tracking is beneficial or hindering (Anderson et al., 2013a; Decruy et al., 2019). In our study, when looking at inter-individual differences in older adults, stronger envelope tracking seems to be a factor that is beneficial for speech understanding in older adults. Specifically, it predicted accuracy in the CT, with which we assessed how well our participants memorized the content of the auditory stimuli. This result is in line with previous findings in young adults, where envelope tracking was positively related to speech-in-noise intelligibility (Ding and Simon, 2013). Possibly, a stronger representation of the envelope in neural activity during the TRF_300_ time window reflects more faithful encoding which in turn results in better recall during the CT trials.

In our view, the debate whether strong envelope tracking in older adults should be considered beneficial or hindering stems at least partly from the heterogeneity of methods employed to measure envelope tracking. From auditory steady-state responses over envelope reconstruction (backward modeling) to TRFs (forward modeling), these methods all highlight different aspects and stages of envelope tracking. While forward modeling allows the investigation of envelope tracking as it unfolds over time, envelope reconstruction/backward modeling has the benefit of providing an actual quantification of reconstruction accuracy. We wish to emphasize that for the case of forward modeling, stronger target speaker envelope tracking seems to be beneficial for speech intelligibility, both in younger (Ding and Simon, 2013) and older adults (the present study). Given that with the FUEL, current theory on effortful listening views selective attention as an essential ability for successful speech-in-noise processing (Pichora-Fuller et al., 2016), our finding that envelope tracking is enhanced in participants with better selective attention provides another cue for a beneficial role of envelope tracking. Furthermore, we showed that stronger envelope tracking in multi-talker babble noise was advantageous even further downstream during speech processing, namely, at the level of memorizing the content of the speech signal. Therefore, the benefits of selective attention and enhanced envelope tracking may well extend beyond object formation and object selection (Shinn-Cunningham, 2008; Shinn-Cunningham and Best, 2008) and remove cognitive load during later stages of speech processing (McCoy et al., 2005).

## Conclusion

This study investigated the influences of EM and IM on speech in noise processing in older adults. There was no additive effect of EM and IM on speech processing, but EM influenced how well participants could follow a target speaker in the presence of background talkers. Further, both EM and IM influenced how well participants could memorize the content of the target speaker’s speech. The amount of speech envelope tracking was affected by IM and modulated by selective attention. Also, the amplitude of a later component of the TRF to the speech envelope, the TRF_300_, was positively related to how well participants could memorize the content of the target speaker’s speech. To summarise, increases in EM and IM both rendered speech-in-noise processing more difficult, and affected speech envelope processing.

## Acknowledgements

This work was supported by grants from the Swiss National Science Foundation (grant no. 172268 to IK and grant no. 169964 to MM), and by the Sonova AG. During the work on their dissertations, IK, LJ, and PN were pre-doctoral fellows of the International Max Planck Research School on the Life Course (LIFE).

